# Whole-genome analysis of antimicrobial-resistant *Escherichia coli* in human gut microbiota reveals its origin and flexibility in transmitting *mcr-1*

**DOI:** 10.1101/2021.02.19.431933

**Authors:** Yichen Ding, Woei-Yuh Saw, Linda Wei Lin Tan, Don Kyin Nwe Moong, Niranjan Nagarajan, Yik Ying Teo, Henning Seedorf

## Abstract

Multidrug resistant (MDR) *Escherichia coli* strains that carry extended-spectrum β-lactamases (ESBLs) or colistin resistance gene *mcr-1* have been identified in the human gut at an increasing incidence worldwide. In this study, we sampled and characterized MDR Enterobacteriaceae from the gut microbiota of healthy Singaporeans and show that the prevalence of ESBL-producing and *mcr*-positive Enterobacteriaceae is 26.6% and 7.3%, respectively. Whole-genome sequencing of 37 *E. coli* isolates identified 25 sequence types and assigned them into six different phylogroups, suggesting that the human intestinal MDR *E. coli* strains are highly diverse. In addition, we found that *E. coli* isolates belonging to phylogroup D, B2 and F carry a higher number of virulence genes, whereas isolates of phylogroup A, B1 and E carry fewer virulence factor genes but are frequent carriers of florfenicol resistance gene *floR* and colistin resistance gene *mcr-1*. Comparison of the seven *mcr-1*-positive *E. coli* isolates revealed that *mcr-1* is carried by conjugative plasmids or embedded in composite transposons, which could potentially mobilize *mcr-1* to other pathogenic Enterobacteriaceae strains or MDR plasmids. Finally, we found that 12 out of the 37 MDR *E. coli* isolates in this study show high similarity to ESBL-producing *E. coli* isolates from raw meats from local markets, suggesting a potential transmission of MDR *E. coli* from meat products to the human gut microbiota. Our findings show diverse antibiotic resistance and virulence profiles of intestinal *E. coli* and call for better countermeasures to block the transmission of MDR *E. coli* via the food chain.

**Importance:** The human gut can harbor both antibiotic resistant and virulent *E. coli* which may subsequently cause infections. In this study, the antibiotic resistance and virulence traits of antibiotic-resistant *E. coli* isolates from human gut microbiota of healthy subjects were investigated. The isolated *E. coli* strains carry a diverse range of antibiotic resistance mechanisms and virulence factor genes, are highly diverse to each other, and are likely to originate from raw meat products from the local markets. Of particular concern are seven *E. coli* isolates which carry colistin resistance gene *mcr-1*. This gene can be mobilized into other pathogens and MDR plasmids, thereby spreading resistance to the last-resort antibiotic colistin. Our findings also suggest that raw meat could serve as important source to transmit MDR bacteria into the human gut microbiota.

## Introduction

*Escherichia coli* is a member of the commensal human gut microbiota, as well as an important pathogen that can cause a diverse range of infections. It was recently proposed that the phylogeny of *E. coli* can be divided into eight phylogroups (A, B1, B2, C, D, E, F, and G), which are usually associated with the lifestyles of the strains ^1^. For instance, commensal and diarrheagenic *E. coli* strains usually belong to phylogroups A and B1, whereas most extraintestinal pathogenic *E. coli* (ExPEC) strains that can cause urinary tract infections, sepsis, and bacteremia in human, belong to phylogroups B2 and D ^2^.

Antibiotic treatment is usually required for chronic and severe *E. coli* infections, especially in immunocompromised patients and the elderly. However, the emergence of multi-drugresistant (MDR) *E. coli* has greatly challenged the effectiveness of antibiotics, and therefore, has become a serious public health concern worldwide. For instance, extended-spectrum β-lactamase (ESBL)-producing *E. coli* showed high-level resistance to penicillin and cephalosporins, which are the first-line drugs to treat *E. coli* infections. It was reported that infections caused by ESBL-producing *E. coli* are associated with significantly higher mortality rates, longer hospitalization periods, and higher medical costs ^3^. Moreover, the emerging *E. coli* strains that carry colistin resistance gene *mcr-1* have disseminated worldwide since their first discovery in China in 2015 ^4, 5^. The increasing incidence of the association between *mcr-1* and ESBL has also been reported, which could further compromise the efficacies of these antibiotics in clinical use ^6^.

The human gut can serve as the reservoir for MDR *E. coli* strains, which could cause life-threatening infections when the person is immunocompromised or has other medical conditions ^7^. For example, fecal carriage of ESBL-producing Enterobacteriaceae increased the risks of post-surgical infections in patients undergone organ transplantation, as well as risks of developing bacteremia in patients with hematological malignancies ^8, 9^. Worse still, some of the MDR *E. coli* lineages residing the human intestine are also ExPEC strains, which possess a broad armament of virulence factors that enable them to cause infections in other parts of the body ^10^. Therefore, it is vital to understand the prevalence and epidemiology of MDR *E. coli* gut colonizers, especially those with ExPEC traits.

In Singapore, both ESBL-producing and *mcr-1*-positive Enterobacteriaceae have been identified in the gut microbiota of the residents, with an estimated prevalence of 26% and 9%, respectively ^11, 12^. In this study, we isolated and sequenced ESBL-producing and *mcr*-positive Enterobacteriaceae isolates from the gut microbiota of a cohort of healthy Singaporeans. We show that *mcr-1* is carried and potentially mobilized by several types of composite transposons and is mainly associated with commensal phenotype. Furthermore, our results also suggest that food is a potential source of MDR *E. coli* in the gut microbiota of Singaporeans.

## Results

### Prevalence of ceftriaxone-resistant and *mcr*-positive Enterobacteriaceae

To study the fecal carriage of ESBL-producing and *mcr*-positive Enterobacteriaceae, we used MacConkey agar supplemented with 2 mg/L ceftriaxone or colistin to screen 109 human fecal samples collected from a cohort of healthy Singaporeans ^13^. In total, we obtained and sequenced 41 non-duplicate Enterobacteriaceae isolates from 35 fecal samples (one to three isolates from each fecal sample). Whole genome sequencing revealed that there were 37 *E. coli*, two *Klebsiella pneumoniae*, one *Citrobacter sedlakii*, and one *Enterobacter cloacae*. Of the 109 fecal samples, the detection rates for ESBL-producing and *mcr*-positive Enterobacteriaceae are 26.6% and 7.3%, respectively, which are comparable in prevalence to previous studies ^11, 12^, whereas carbapenemase-producing Enterobacteriaceae remained at a much lower detection rate of 2.8%.

The acquired antibiotic resistance genes and resistance phenotypes of the four non-*E. coli* isolates are shown in Table S1. Briefly, *E. cloacae* 97COLEN showed colistin MIC of > 16 mg/L, which can be explained by the acquisition of *mcr-10*, a novel *mcr* variant first identified in clinical isolate of *Enterobacter roggenkampii* ^14^. While the other three strains showed ceftriaxone MIC of > 64 mg/L, and the resistance mechanisms varied depending on their taxonomy: *C. sedlakii* 18CS produced both carbapenemase *bla*_CMY-2_ and ESBL *bla*_SED-1_, *K. pneumoniae* 102KP produced CTX-M-type ESBL *bla*_CTX-M-14_, and *K. pneumoniae* 25KP produced both *bla*_CTX-M-24_ and SHV-type ESBL *bla*_SHV-75_ (Table S1). We further focused on analysing the 37 *E. coli* isolates, as they represent the majority of the isolates collected in this study.

### Phylogenetic diversity of *E. coli* isolates

In total, 32 *E. coli* isolates showed a ceftriaxone MIC of ≥ 64 mg/L (Table S2). Whole genome sequencing revealed that 30 ceftriaxone-resistant *E. coli* isolates produced CTX-M-type ESBLs, whereas two isolates produced CMY-type carbapenemase (Figure 1). The most commonly identified ESBL is CTX-M-15 (n=13), followed by CTX-M-55 (n=6), CTX-M-14 (n=4), CTX-M-27 (n=4), CTX-M-65 (n=3), and OXA-1 (n=1, this isolate also produces CTX-M-15). Of note, two CTX-M-producing strains, 40EC and 94EC also carried *mcr-1*, making them resistant to both third-generation cephalosporins and colistin (Figure1, Table S2). We further isolated five *mcr-1* positive *E. coli*, all of which were negative for ESBL or carbapenemase, but produced penicillinase *bla*_TEM-1B_ (Figure 1). These five isolates showed resistance to ampicillin and colistin, whereas they remained susceptible to ceftriaxone (Table S2).

**Figure 1.**
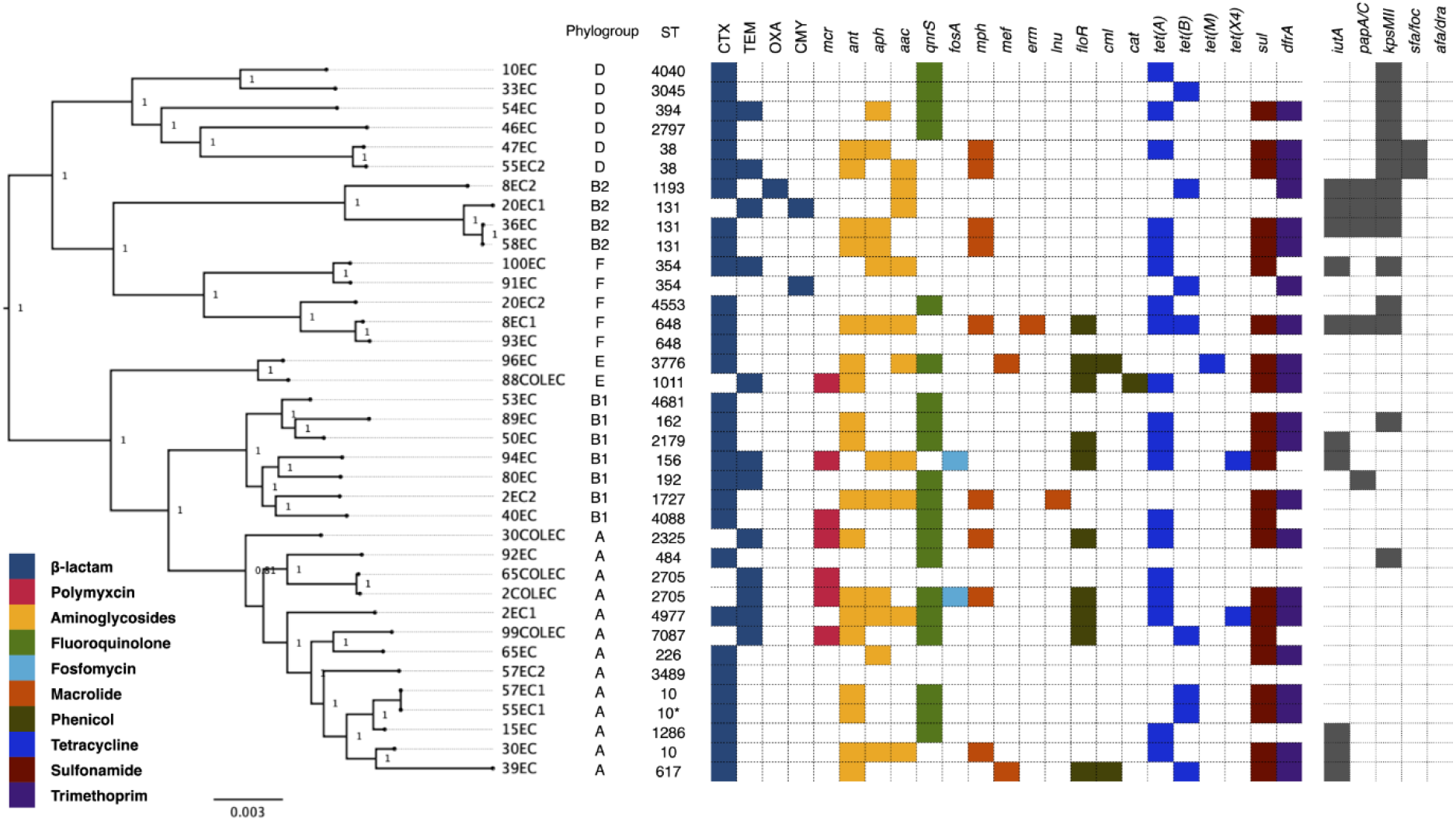
Comparative genomic analysis of ESBL/carbapenemase-producing and *mcr-1-* positive *E. coli* isolates. The maximum-likelihood phylogenetic tree was constructed using the detected variant sites of the core-genome alignment, and the clade confidence was estimated with SH-like supporting values. Colored blocks indicate the presence of acquired antibiotic resistance genes, whereas grey blocks indicate the presence of five selected ExPEC marker genes.

The core-genome phylogenetic tree of the 37 *E. coli* isolates indicates two main clusters, of which 15 isolates belonging to phylogroup D (n = 6), B2 (n = 4) and F (n = 5) formed cluster 1, whereas 22 isolates belonging to phylogroup E (n = 2), B1 (n = 7) and A (n = 13) formed cluster 2 (Figure 1). Multilocus sequence typing assigned 36 isolates into 29 known sequence types (STs), with one ST (ST131) comprised of three isolates, four STs (ST38, ST354, ST648, ST2705, and ST10) comprised of two isolates, and 24 STs comprised of one isolate. The lack of a dominant ST among the 37 isolates suggested that clonal expansion of ESBL-producing or *mcr-1*-positive *E. coli* in the gut microbiota of the 109 subjects is unlikely.

### Antibiotic resistance and virulence factor gene profiling

We further analyzed the acquired antibiotic resistance and virulence factor (VF) gene profiles of the 37 isolates. Other than beta-lactamases, the most commonly identified antibiotic resistance mechanisms are aminoglycosides (*ant, aph*, and *aac*), tetracycline (*tet*(*A*) and *tet*(*B*)), sulfonamide (*sul*), and trimethoprim (*dfrA*), all of which are detected in isolates of both clusters (Figure 1). Interestingly, all the seven *mcr-1* positive isolates are located in cluster 2, including one phylogroup E isolate, two phylogroup B1 isolate, and four phylogroup A isolate (Figure 1). Similarly, nine out of ten isolates carrying chloramphenicol and florfenicol resistance gene *floR* are located in cluster 2, whereas only one *floR*-positive isolate is located in cluster 1 (Figure 1).

Prediction of VF gene showed that cluster 1 isolates carried an average of 13.4 ± 4.2 VF genes, which is significantly higher than cluster 2 isolates with an average of 7.0 ± 4.5 VF genes (Figure S1, P < 0.0001). We did not identify any VFs that are associated with diarrheagenic *E. coli*, which is consistent with the fact that all subjects had no signs of diarrhea upon fecal sampling. However, seven isolates were positive for at-least two ExPEC marker genes, suggesting that they are ExPEC strains (Figure 1) ^2^. All seven ExPEC isolates are found in cluster 1, with two phylogroup D isolates, three phylogroup B2 isolates, and two phylogroup F isolates (Figure 1). Several other VF genes were also more commonly found in cluster 1 isolates, including *air* (enteroaggregative immunoglobulin repeat protein), *chuA* (outer membrane heme receptor), *eilA* (Salmonella HilA homolog), *fyuA* (siderophore receptor), *irp2* (non-ribosomal peptide synthase), *iss* (serum survival), *kpsE* (capsule polysaccharide export), *sitA* (iron transport), *usp* (uropathogenic specific protein), and *yfcA* (fimbriae protein) (Figure S1). By contrast, the long polar fimbriae gene *lpfA* was mainly identified in phylogroup F and B1 isolates, which belong to cluster 1 and cluster 2, respectively. The carriage of *lpfA* was suggested to contribute to their gut colonization ^15^.

Taken together, our analysis clearly show that ESBL-producing and *mcr-1*-positive *E. coli* isolates in this study can be divided into two clusters based on their core-genome phylogeny (Figure 1). Cluster 1 isolates are associated with a more virulent phenotype and include all seven ExPEC strains, whereas cluster 2 isolates have fewer VF genes but contain most of the isolates that carried *floR* and *mcr-1*, which are responsible for resistance to florfenicol and colistin, respectively (Figure 1).

### Transmission of MDR *E. coli* between human and raw meat

In a recent study, ESBL-producing and *mcr*-positive *E. coli* have been isolated and characterized from retail raw meat in Singapore ^16^. This made us wonder if similarity between ESBL-producing *E. coli* strains from raw meat and human subjects could be observed. We therefore constructed a core-genome phylogenetic tree consisting of 37 *E. coli* isolates sequenced in this study and 225 ESBL-producing *E. coli* isolates from raw meat from local markets ^16^. Interestingly, we found that 32.4% of the *E. coli* isolates (12/37) characterized in our study are associated with at-least one raw meat isolate as indicated by phylogenetic and MLST analysis (Figure 2), strongly suggesting the possible transmission of MDR *E. coli* from raw meat to the human gut microbiota. Of the 10 clades that contained both human fecal isolates and raw meat isolates of the same ST, the ST156 clade contained six isolates from raw chicken meat, one isolate from raw beef meat and one isolate from human feces (94EC), all of which carried CTX-M-65 type ESBL, *mcr-1* and the recently identified tigecycline resistance gene *tet*(X4) (Figure 2). The carriage of resistance mechanisms to three classes of last-resort antibiotics could make the further spread of this clone a serious public health concern.

**Figure 2.**
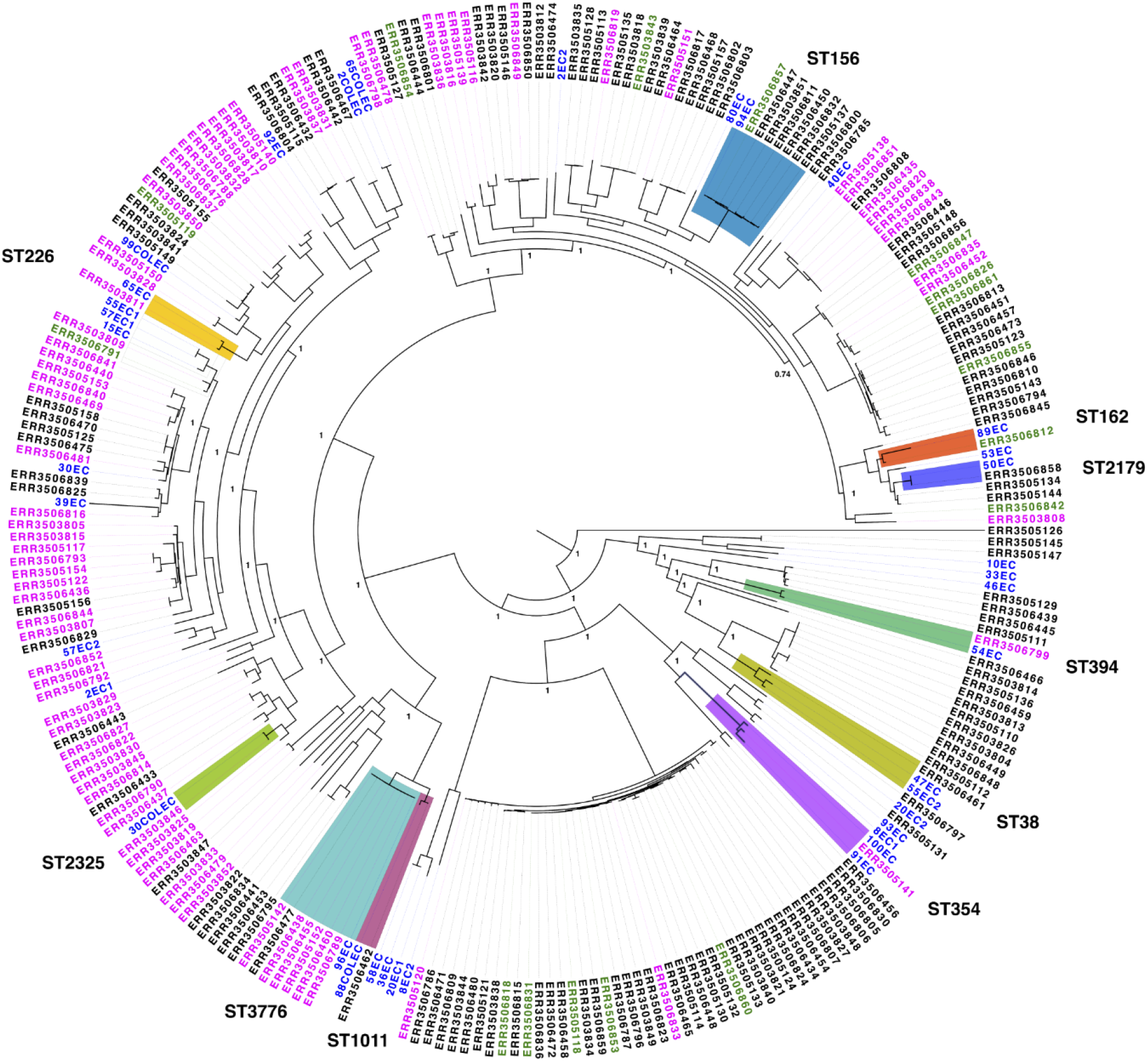
Phylogenetic analysis of 37 *E. coli* isolates from human feces and 225 ESBL-producing *E. coli* isolates from retail raw meats. The maximum-likelihood phylogenetic tree was constructed based on the core-genome SNPs of 37 *E. coli* isolates in this study, and 225 ESBL-producing *E. coli* isolates from a previous study ^16^. The source of the isolates is indicated by the color of the label: blue (isolates from human fecal samples, this study), black (isolates from raw chicken meat), magenta (isolates from raw pork meat), and green (isolates from raw beef meat). The clades highlighted in colors fulfills three criteria: 1) it contains isolates from both human feces and raw meat; 2) the isolates have the same ST; 3) the genomes of the isolates in the clade share an ANI similarity of at-least 99.4%. The clade confidence was estimated with SH-like supporting values, which are shown for the major clades in the figure. The genomes of raw meat isolates were assembled from sequencing data deposited at the European Nucleotide Archive (https://www.ebi.ac.uk/ena) under accession number PRJEB34067.

### Genetic context of *mcr-1*

Colistin resistance gene *mcr-1* has been shown to be mobilized by *ISApl1* and found on both conjugative plasmids and the chromosomes of Enterobacteriaceae ^17^. We identified four different types of *mcr-1*-containing cassettes in the seven *mcr-1*-positive *E. coli* isolates, suggesting the highly diversified genetic context of *mcr-1* (Figure 3). Five *mcr*-1 insertional events happened on the *E. coli* chromosomes, whereas three isolates carried plasmids that contain *mcr-1*. The comparison of the four types of *mcr*-1-containing cassettes is shown in Figure 3.

**Figure 3.**
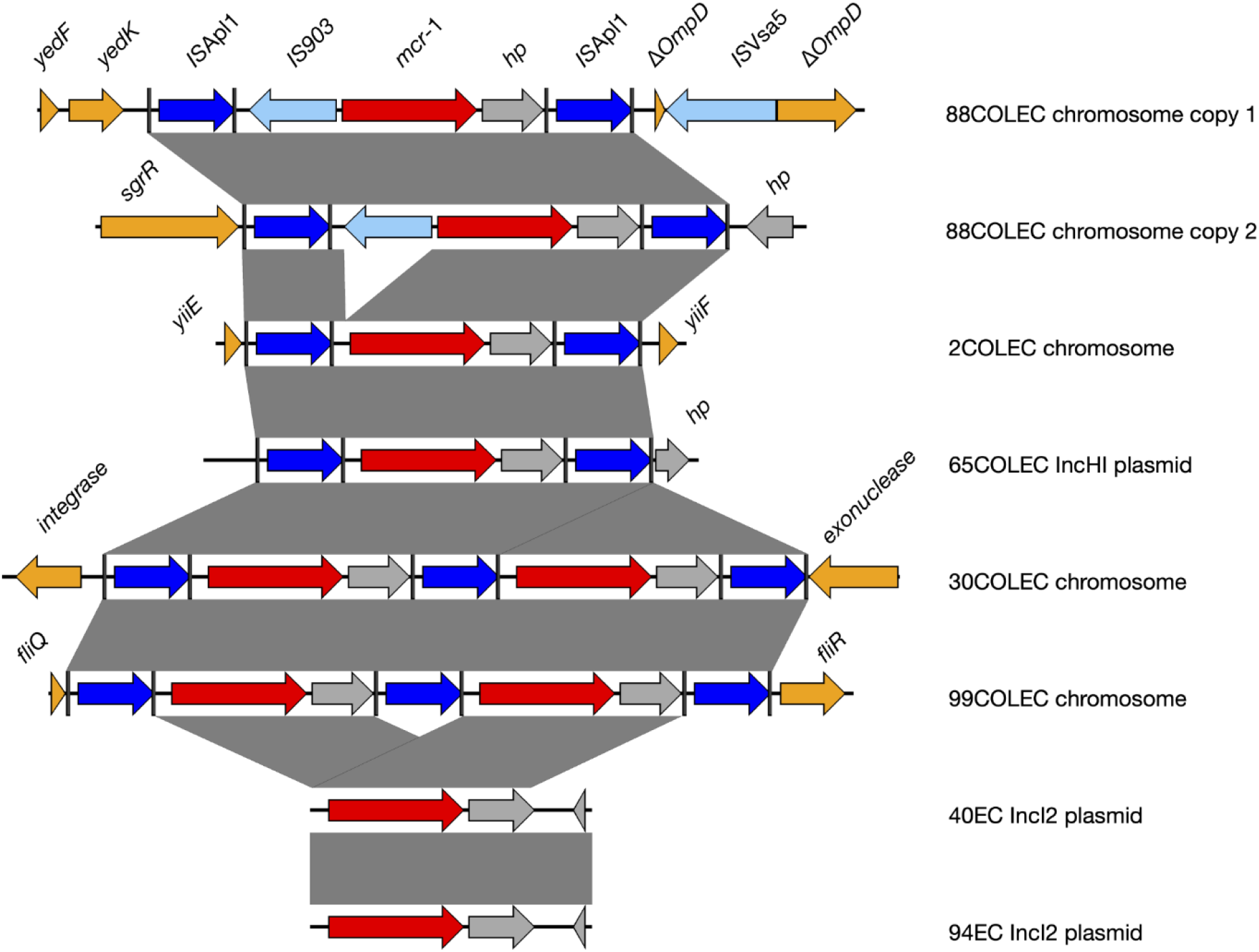
Sequence alignment of *mcr-1*-containing cassettes. Arrow indicates open reading frame and the direction of transcription. IS*Apl1* is indicated by blue arrow, *mcr-1* is indicated by red arrow, hypothetical gene is indicated by grey arrow, and intrinsic genes of the plasmid backbone or chromosome are indicated by orange arrows. Grey shadings connect regions of > 99% similarity. Black vertical lines denote left and right inverted repeats of IS*Apl1*.

Firstly, the *ISApl1-mcr-1-hp-ISApl1* cassette was identified in isolates 2COLEC and 65COLEC, both of which belong to ST2705 (Figure 1 and 3). This structure was described as *Tn6330,* a composite transposon that can generate circular intermediate harboring *mcr-1* and IS*Apl1* ^18^. Interestingly, although the genomes of 2COLEC and 65COLEC genomes are highly similar, *Tn6330* was inserted in the chromosome of 2COLEC, but was inserted in an IncHI plasmid in 65COLEC (Figure 3). Secondly, the *ISApl1-mcr-1-hp-ISApl1-mcr-1-hp-ISApl1* cassette resulted from tandem duplication of Tn*6330* and was identified in the chromosomes of 30COLEC (between *fliQ* and *fliR*) and 99COLEC (between a phage integrase gene and a gene encoding exonuclease) (Figure 3). Tandem duplication of Tn*6330* has not been described before, while a previous study has identified a triplicated Tn*6330* structure in a clinical *E. coli* isolate ^19^. Thirdly, the *ISApl1-IS903-mcr-1-hp-ISApl1* cassette with an additional IS*903* element inserted between the upstream *ISApl1* and *mcr-1* was identified in isolate 88COLEC. We found two copies of *ISApl1-IS903-mcr-1-hp-ISApl1* cassettes in 88COLEC chromosome: one was located between *yedK* and a truncated *ompD* gene, whereas the other one was located between *sgrR* and a hypothetical gene (Figure 3). In all these three types of cassettes, *mcr-1* was bound by an upstream and a downstream IS*Apl1* with intact left and right inverted repeats (Figure 3), thereby forming composite transposon structures that can generate circular intermediates by a cut-and-paste mechanism ^18, 20^. Consistently, PCR using reverse primers targeting *mcr-1* confirmed the presence of circular intermediates in all the five isolates (Figure S2), suggesting that they can mobilize *mcr-1* into the backbone of other MDR plasmids to associate *mcr-1* with other antibiotic resistance mechanisms. Lastly, *mcr-1* was located on IncI plasmids p40EC-4 and p94EC-3 of 40EC and 94EC, respectively, and have lost both upstream and downstream IS*Apl1*. p40EC-4 and p94EC-3 share a similarity of 99.68% and are highly similar to other *mcr-1* carrying IncI plasmids reported in previous studies ^18, 21^. The loss of IS*Apl1* by *mcr-1* was suggested to contribute to the stability of the plasmid replicon, causing the mobilization of *mcr-1* solely to rely on the conjugative transfer of the plasmid ^22^.

## Discussion

In this study, we sequenced the genomes of ceftriaxone-resistant and *mcr*-positive Enterobacteriaceae isolated from fecal samples of a cohort of healthy Singaporeans. We report that the fecal carriage rate of ESBL-positive *mcr*-positive Enterobacteriaceae are 26.6% and 7.3%, respectively. The most commonly isolated ESBL-producing Enterobacteriaceae is *E. coli* that produces CTX-M-type ESBL (n = 30), whereas the most commonly isolated *mcr*-positive Enterobacteriaceae is *mcr-1*-positive *E. coli* (n = 7). These findings are consistent with previous studies, which reported that the fecal carriage rate of ESBL-producing Enterobacteriaceae is 26.2% in a healthy community, whereas the fecal carriage rate of *mcr-1*-positive *E. coli* is estimated to be 9% in diarrheal patients in Singapore ^11, 12^. Although our isolates belong to a diverse range of STs, we found ST131 to be more common among the ceftriaxone-resistant *E. coli* than other STs (10%, 3/30). Of the three ST131 isolates, 20EC1 and 36EC are ExPEC strains and produced carbapenemase CMY and CTX-M-type ESBL, whereas 36EC is not ExPEC and produced CTX-M-type ESBL (Figure 1). This suggested that the ceftriaxone-resistant *E. coli* ST131 isolated in our study is associated with versatile resistance mechanisms and virulence phenotypes, which is consent with the global and local dissemination of ST131 strains as colonizer and causative agent for infections ^10, 11, 23^.

ExPEC strains can colonize the human gut, where they can later disseminate and cause extraintestinal infections ^24^. We identified seven ExPEC isolates, including two ST131 isolates, two ST38 isolates, one ST1193 isolate, one ST354 isolate, and one ST648 isolate (Figure 1). All the five STs were among the top 20 ExPEC STs that account for 85% of the ExPEC isolates across 169 studies, as revealed by a meta-analysis ^10^. Interestingly, the seven ExPEC strains identified here seemed to be negatively associated with the fluoroquinolone resistance gene *qnrS* (Figure 1), suggesting that fluoroquinolone could be considered for infections caused by ESBL/carbapenemase-producing ExPEC strains locally.

Our phylogenetic analysis indicated that 10 of the 37 MDR *E. coli* isolates from human gut microbiota share high similarity with isolates from raw meat products, suggesting that raw meat could serve as one of the major sources for MDR *E. coli* in the gut microbiota of Singaporeans (Figure 2). This finding is consistent with previous studies conducted in Europe ^25, 26^. Our previous study revealed that the ESBL-producing, *mcr-1* positive and *tet*(X4)-positive strain 94EC is the dominant Enterobacteriaceae strain in the gut microbiota of a subject, suggesting that it is able to establish an ecological niche in human gut microbiota and may also be associated with higher risks to spread in the community ^27^. Consistently, seven isolates from raw meat collected between 2017 and 2018 ^16^ shared the same ST and antibiotic resistance genes with 94EC (Figure 2), which was isolated from a fecal sample collected in 2018 ^27^. This evidence suggests a possible silent outbreak of the MDR ST156 clone in the community during that time, while its further dissemination in the community should be closely monitored.

We found that *mcr-1* is strongly associated with resistance mechanisms to beta-lactams: two *mcr-1* positive *E. coli* also produces CTX-M-type ESBL, whereas the other five *E. coli* produces penicillinase *bla*_TEM-1B_. A previous study conducted in Guangzhou China has shown a rapid increase of association between *mcr-1* and cefotaxime resistance between 2014 and 2016 ^6^. In our study, the mobilization of *mcr-1* gene is mostly related to IS*Apl1*-mediated transposition (Figure 3 and S2). This process can facilitate the emergence of novel MDR Enterobacteriaceae strains with colistin resistance. Therefore, surveillance of local *mcr-1* dissemination should be conducted. In addition, the two IncI plasmids p40EC-4 and p94EC-3 are highly similar to IncI type *mcr-1*-carrying plasmids reported previously in China ^18, 21^, suggesting a potential direct epidemiological linkage.

In conclusion, we found that the ceftriaxone-resistant and colistin-resistant *E. coli* fecal isolates from Singaporeans belong to a diverse range of phylogroups and ST, contained subgroups with unique virulence and antibiotic resistance profiles, and may originate from raw meat from local markets. Furthermore, we also found *mcr-1* to be located on highly actively composite transposons, which may contribute to new combinations of MDR plasmids and pathogenic strains. Our findings strongly suggest MDR *E. coli* strains could transmit from food to human and call for better countermeasures to be implemented.

## Methods

### Strain isolation and characterization

Fecal samples from 109 individuals of the Singapore Integrative Omics Study were collected in 2018 using BioCollector kit (Biocollective) as per manufacturer’s instructions ^28^. Fecal samples were suspended in LB broth and incubated at 37 °C with 200 rpm shaking for three hours, before being spread onto MacConkey agar plates supplemented with 2 mg/L ceftriaxone or 2 mg/L colistin. For each observed colony morphology type on the selective plate, three colonies were initially picked and assessed by Random Amplification of Polymorphic DNA analysis using M13-core primer to verify possible duplication ^29^. The verified non-duplicate colonies were subjected to 16S rRNA gene sequencing using 27F and 1492R primers ^30^. Species that are intrinsically resistant to colistin, including *Proteus mirabilis, Morganella morganii, Hafnia paralvei*, and *Providencia rettgeri* were excluded from whole-genome sequencing.

### Genome sequencing and assembly

Genomic DNA was purified using Qiagen Genomic-tip 20/G (Qiagen) as per manufacturer’s instructions. Whole-genome sequencing was first performed using Illumina Novaseq that generated 150 bp paired-end reads, which were assembled into draft genomes by SPAdes v3.11.1 with --careful option on and k-mer sizes of 21, 33, 55, 77, 99, 127 ^31^. For complete genome assembly, the genomic DNA was further sequenced on MinION (Oxford Nanopore Technologies) using R9.4.1 flow cell and Rapid Barcoding Kit as per manufacturer’s instructions. Sequencing reads from both MinION and Illumina were used for hybrid assembly by Unicycler v0.4.8 ^32^.

### Comparative genomic analysis

Phylogenetic tree of *E. coli* genomes was constructed using the Harvest Suite ^33^. Briefly, the core-genome was aligned by Parsnp v1.1.2 ^33^, and the variant sites in the core-genome were used to infer phylogenetic relationship using approximate maximum likelihood algorithm by FastTree 2 integrated in Parsnp ^34^. The branch length of the phylogenetic tree represents substitutions per core-genome site, and the clade confidence was estimated by SH-like supporting values ^33^. Gene prediction and annotation was performed by the RAST server ^35^, acquired antibiotic resistance genes were predicted by ResFinder v4.1 ^36^, insertional sequences were identified by ISFinder ^37^, virulence factor genes were predicted by Virulence Finder v2.0 ^38^, *in silico* multilocus sequence typing was performed by MLST server ^39^, and phylogroup was identified by Clermontyping server ^1, 40^. The phylogenetic tree was visualized using FigTree v1.4.4, and comparative analysis of sequences was performed using EasyFig v2.2.2.^41^. Average nucleotide identity calculation was performed by ANI calculator ^42^.

### Antibiotic susceptibility test

The minimal inhibitory concentrations of ampicillin, ceftriaxone and colistin was performed according to CLSI guidelines by broth microdilution in accordance to CLSI guidelines ^43^. Cation-adjusted Mueller Hinton broth was used for MIC experiment, with an incubation time of 18 hours at 37 °C.

### PCR detection of circular *mcr-1* intermediates

The circular *mcr-1* intermediates are detected by PCR using primers MCR1-RC-F (ACGCACAGCAATGCCTATGA) and MCR1-R (CTTGGTCGGTCTGTAGGG) as previously described ^18^. One μl of bacterial overnight culture was used as template for PCR.

### Ethics

This study was conducted in compliance with the Declaration of Helsinki and national and institutional standards. The collection of human faecal samples for this study was approved under the National University of Singapore IRB code H-17-026.

---

**Figure S1** Virulence gene profiling of *E. coli isolates.* Blue blocks indicate the presence of virulence genes in the genome.

**Figure S2** Detection for circular intermediates containing *mcr-1*. M: 1kb marker. 1: 40EC, 2: 94EC, 3: 2COLEC, 4: 30COLEC, 5: 65COLEC, 6: 97COLEN, 7: 99COLEC, 8: 88COLEC. The expected size of PCR products in lane 3, 4, 5 and 7 is 2672 bp, whereas lane 8 shows a larger PCR product of 3728 bp due to the insertion of an IS*903* element (Figure 3).

## References

1. Clermont O, Dixit OVA, Vangchhia B et al. Characterization and rapid identification of phylogroup G in *Escherichia coli*, a lineage with high virulence and antibiotic resistance potential. Environ Microbiol 2019; 21: 3107–17.

2. Johnson JR, Murray AC, Gajewski A et al. Isolation and molecular characterization of nalidixic acid-resistant extraintestinal pathogenic *Escherichia coli* from retail chicken products. Antimicrob Agents Chemother 2003; 47: 2161–8.

3. Schwaber MJ, Navon-Venezia S, Kaye KS et al. Clinical and economic impact of bacteremia with extended-spectrum-beta-lactamase-producing Enterobacteriaceae. Antimicrob Agents Chemother 2006; 50: 1257–62.

4. Liu YY, Wang Y, Walsh TR et al. Emergence of plasmid-mediated colistin resistance mechanism MCR-1 in animals and human beings in China: a microbiological and molecular biological study. Lancet Infect Dis 2016; 16: 161–8.

5. Ling Z, Yin W, Shen Z et al. Epidemiology of mobile colistin resistance genes *mcr-1* to *mcr-9*. J Antimicrob Chemother 2020; 75: 3087–95.

6. Zhong LL, Phan HTT, Shen C et al. High Rates of Human Fecal Carriage of *mcr-1*-Positive Multidrug-Resistant Enterobacteriaceae Emerge in China in Association With Successful Plasmid Families. Clin Infect Dis 2018; 66: 676–85.

7. Sommer MOA, Dantas G, Church GM. Functional characterization of the antibiotic resistance reservoir in the human microflora. Science 2009; 325: 1128–31.

8. Bert F, Larroque B, Paugam-Burtz C et al. Pretransplant fecal carriage of extended-spectrum beta-lactamase-producing Enterobacteriaceae and infection after liver transplant, France. Emerg Infect Dis 2012; 18: 908–16.

9. Cornejo-Juarez P, Suarez-Cuenca JA, Volkow-Fernandez P et al. Fecal ESBL *Escherichia coli* carriage as a risk factor for bacteremia in patients with hematological malignancies. Support Care Cancer 2016; 24: 253–9.

10. Manges AR, Geum HM, Guo A et al. Global Extraintestinal Pathogenic *Escherichia coli* (ExPEC) Lineages. Clin Microbiol Rev 2019; 32.

11. Mo Y, Seah I, Lye PSP et al. Relating knowledge, attitude and practice of antibiotic use to extended-spectrum beta-lactamase-producing Enterobacteriaceae carriage: results of a cross-sectional community survey. BMJ Open 2019; 9: e023859.

12. La MV, Lee B, Hong BZM et al. Prevalence and antibiotic susceptibility of colistin-resistance gene (*mcr-1*) positive Enterobacteriaceae in stool specimens of patients attending a tertiary care hospital in Singapore. Int J Infect Dis 2019; 85: 124–6.

13. Saw WY, Tantoso E, Begum H et al. Establishing multiple omics baselines for three Southeast Asian populations in the Singapore Integrative Omics Study. Nat Commun 2017; 8: 653.

14. Wang C, Feng Y, Liu L et al. Identification of novel mobile colistin resistance gene *mcr-10*. Emerg Microbes Infect 2020; 9: 508–16.

15. Torres AG, Giron JA, Perna NT et al. Identification and characterization of lpfABCC’DE, a fimbrial operon of enterohemorrhagic *Escherichia coli* O157:H7. Infect Immun 2002; 70: 5416–27.

16. Guo S, Aung KT, Leekitcharoenphon P et al. Prevalence and genomic analysis of ESBL-producing *Escherichia coli* in retail raw meats in Singapore. J Antimicrob Chemother 2020.

17. Wang R, van Dorp L, Shaw LP et al. The global distribution and spread of the mobilized colistin resistance gene *mcr-1*. NatCommun 2018; 9: 1179.

18. Li R, Xie M, Zhang J et al. Genetic characterization of mcr-1-bearing plasmids to depict molecular mechanisms underlying dissemination of the colistin resistance determinant. J Antimicrob Chemother 2017; 72: 393–401.

19. Yu CY, Ang GY, Chong TM et al. Complete genome sequencing revealed novel genetic contexts of the *mcr-1* gene in *Escherichia coli* strains. J Antimicrob Chemother 2017; 72: 12535.

20. Li R, Xie M, Lv J et al. Complete genetic analysis of plasmids carrying *mcr-1* and other resistance genes in an *Escherichia coli* isolate of animal origin. J Antimicrob Chemother 2017; 72: 696–9.

21. Li XP, Sun RY, Song JQ et al. Within-host heterogeneity and flexibility of *mcr-1* transmission in chicken gut. Int J Antimicrob Agents 2020; 55: 105806.

22. Snesrud E, He S, Chandler M et al. A Model for Transposition of the Colistin Resistance Gene *mcr-1* by IS*Apl1*. Antimicrob Agents Chemother 2016; 60: 6973–6.

23. Harris PNA, Ben Zakour NL, Roberts LW et al. Whole genome analysis of cephalosporin-resistant Escherichia coli from bloodstream infections in Australia, New Zealand and Singapore: high prevalence of CMY-2 producers and ST131 carrying *bla*_CTX-M-15_ and *bla*_CTX-M-27_. J Antimicrob Chemother 2018; 73: 634–42.

24. Yamamoto S, Tsukamoto T, Terai A et al. Genetic evidence supporting the fecal-perineal-urethral hypothesis in cystitis caused by *Escherichia coli*. J Urol 1997; 157: 1127–9.

25. Kluytmans JA, Overdevest IT, Willemsen I et al. Extended-spectrum beta-lactamase-producing *Escherichia coli* from retail chicken meat and humans: comparison of strains, plasmids, resistance genes, and virulence factors. Clin Infect Dis 2013; 56: 478–87.

26. Leverstein-van Hall MA, Dierikx CM, Cohen Stuart J et al. Dutch patients, retail chicken meat and poultry share the same ESBL genes, plasmids and strains. Clin Microbiol Infect 2011; 17: 873–80.

27. Ding Y, Saw WY, Tan LWL et al. Emergence of tigecycline- and eravacycline-resistant Tet(X4)-producing Enterobacteriaceae in the gut microbiota of healthy Singaporeans. J Antimicrob Chemother 2020; 75: 3480–4.

28. Saw W-Y, Tantoso E, Begum H et al. Establishing multiple omics baselines for three Southeast Asian populations in the Singapore Integrative Omics Study. Nature communications 2017; 8: 1–11.

29. Vogel L, van Oorschot E, Maas HM et al. Epidemiologic typing of *Escherichia coli* using RAPD analysis, ribotyping and serotyping. Clin Microbiol Infect 2000; 6: 82–7.

30. Jiang H, Dong H, Zhang G et al. Microbial diversity in water and sediment of Lake Chaka, an athalassohaline lake in northwestern China. Appl Environ Microbiol 2006; 72: 3832–45.

31. Bankevich A, Nurk S, Antipov D et al. SPAdes: a new genome assembly algorithm and its applications to single-cell sequencing. J Comput Biol 2012; 19: 455–77.

32. Wick RR, Judd LM, Gorrie CL et al. Unicycler: resolving bacterial genome assemblies from short and long sequencing reads. PLoS computational biology 2017; 13: e1005595.

33. Treangen TJ, Ondov BD, Koren S et al. The Harvest suite for rapid core-genome alignment and visualization of thousands of intraspecific microbial genomes. Genome Biol 2014; 15: 524.

34. Price MN, Dehal PS, Arkin AP. FastTree 2--approximately maximum-likelihood trees for large alignments. PLoS One 2010; 5: e9490.

35. Aziz RK, Bartels D, Best AA et al. The RAST Server: rapid annotations using subsystems technology. BMCgenomics 2008; 9: 75.

36. Bortolaia V, Kaas RS, Ruppe E et al. ResFinder 4.0 for predictions of phenotypes from genotypes. J Antimicrob Chemother 2020; 75: 3491–500.

37. Siguier P, Perochon J, Lestrade L et al. ISfinder: the reference centre for bacterial insertion sequences. Nucleic Acids Res 2006; 34: D32–6.

38. Joensen KG, Scheutz F, Lund O et al. Real-time whole-genome sequencing for routine typing, surveillance, and outbreak detection of verotoxigenic *Escherichia coli*. J Clin Microbiol 2014; 52: 1501–10.

39. Larsen MV, Cosentino S, Rasmussen S et al. Multilocus sequence typing of total-genome-sequenced bacteria. J Clin Microbiol 2012; 50: 1355–61.

40. Beghain J, Bridier-Nahmias A, Le Nagard H et al. ClermonTyping: an easy-to-use and accurate in silico method for Escherichia genus strain phylotyping. Microb Genom 2018; 4.

41. Sullivan MJ, Petty NK, Beatson SA. Easyfig: a genome comparison visualizer. Bioinformatics 2011; 27: 1009–10.

42. Yoon SH, Ha SM, Lim J et al. A large-scale evaluation of algorithms to calculate average nucleotide identity. Antonie Van Leeuwenhoek 2017; 110: 1281–6.

43. CLSI. Performance Standards for Antimicrobial Susceptibility Testing—Thirty-Third Edition: M100. 2020.

